# A quick and easy way to estimate entropy and mutual information for neuroscience

**DOI:** 10.1101/2020.08.04.236174

**Authors:** Mickael Zbili, Sylvain Rama

**Affiliations:** Lyon Neuroscience Research Center, INSERM U1028-CNRS UMR 5292-Université Claude Bernard Lyon1 - Lyon, France; Laboratory of Synaptic Imaging, Department of Clinical and Experimental Epilepsy, UCL Queen Square Institute of Neurology, University College London - London, United Kingdom

**Keywords:** Entropy, Mutual Information, PNG, DEFLATE, Rastergram, Lossless compression, Place Fields

## Abstract

Calculations of entropy of a signal or mutual information between two variables are valuable analytical tools in the field of neuroscience. They can be applied to all types of data, capture nonlinear interactions and are model independent. Yet the limited size and number of recordings one can collect in a series of experiments makes their calculation highly prone to sampling bias. Mathematical methods to overcome this so-called “sampling disaster” exist, but require significant expertise, great time and computational costs. As such, there is a need for a simple, unbiased and computationally efficient tool for estimating the level of entropy and mutual information. In this paper, we propose that application of entropy-encoding compression algorithms widely used in text and image compression fulfill these requirements. By simply saving the signal in PNG picture format and measuring the size of the file on the hard drive, we can estimate entropy changes through different conditions. Furthermore, with some simple modifications of the PNG file, we can also estimate the evolution of mutual information between a stimulus and the observed responses through different conditions. We first demonstrate the applicability of this method using white-noise-like signals. Then, while this method can be used in all kind of experimental conditions, we provide examples of its application in patch-clamp recordings, detection of place cells and histological data. Although this method does not give an absolute value of entropy or mutual information, it is mathematically correct, and its simplicity and broad use make it a powerful tool for their estimation through experiments.

## Introduction

Entropy is the major component of information theory, conceptualized by Shannon in 1948 (Shannon, 1948). It is a dimensionless quantity representing uncertainty about the state of a continuous or discrete system or a collection of data. It is highly versatile as it applies to many different types of data, it can capture nonlinear interactions, and is model independent (Cover and Thomas, 2006). It has been widely used in the field of neurosciences, see (Borst and Theunissen, 1999; Piasini and Panzeri, 2019; Timme and Lapish, 2018) for a more complete review of work; for example in the field of synaptic transmission (London et al., 2002), information rate of Action Potentials (APs) trains (Bialek et al., 1991; Juusola and de Polavieja, 2003; Street, 2020) or connectivity studies (Ito et al., 2011; Vicente et al., 2011).

However, estimating the entropy of a signal can be a daunting task. The entropy *H* of a signal X is calculated with the well-known Shannon’s formula:

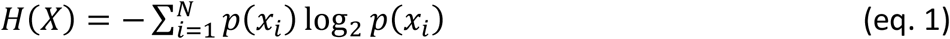

Where *p(x*_*i*_) is the probability that the signal will take the x_i_ configuration among all the configurations (x_1_, x_2_, x_3_,…, x_N_) of the signal. It is considered that if *p*(*x*_*i*_) =0, then *p*(*x*_*i*_)log_2_ *p*(*x*_*i*_)=0, as lim_*x* →0_ *x*(log_2_ *x*) =0. And using a base 2 logarithm, entropy will be expressed in bits (Cover and Thomas, 2006; Shannon, 1948).

However, correctly estimating a probability distribution works only if each configuration happens many times. And by definition, one cannot know beforehand the number of needed experiments. This recording bias is even amplified by the fact that without making assumptions, there is no way to determine the relevant quantization and sampling of the data, i.e. in which probability space the entropy must be calculated. The same recordings could be divided in any quantization bins and sampled by any interval, all giving different probability distributions and thus different entropy values.

As an example, let us consider the chain of characters A=“04050405”. It is unchanged with a quantization range *v* of 6, but will become “01010101” with a quantization range *v* of 2. If we now combine those characters with a bin *T* of 1, this will give a probability distribution of: p(0) = 0.5, p(4) = p(5) = 0.25 in the first scenario (*v* = 6) and: *p(0)* = *p(1)* = 0.5 in the second scenario (*v* = 2). We thus obtain different entropy values for these two probability spaces: *H*^*v*=*6,T*=1^1.5 and *H*^*v*=*2,T*=1^ =1. Now, if we take a combination bin of *T* = 2 we obtain p(04) = p(05) = 0.5 for *v* = 6 and p(01) = 1 for *v* = 2. The calculated entropies thus are: *H*^*v*=*6,T*=2^=1 and *H*^*v*=*2,T*=2^=0 In this example we could take any different of *v* and *T* and obtain different values for the entropy.

Without making assumptions on the data, there is no way to determine which probability space and which value of entropy is the correct one. Therefore, quantization range and combination bins are crucial to determine the entropy of a signal. In an ideal case, we would need a method able to correct for this sample bias without making assumptions about the signal, and exploring every possible probability space meaning for any length of acquisition, any quantization range *v* and any *T* combination bins of the recorded data.

Thankfully there are multiple ways to use the Shannon’s formula (eq. 1) and compensate for this bias, but none of them can be called trivial. There are for example the quadratic extrapolations method (Juusola and de Polavieja, 2003; de Polavieja et al., 2005; Strong et al., 1998), the Panzeri-Treves Bayesian estimation (Panzeri and Treves, 1996), the Best Universal Bound estimation (Paninski, 2003), the Nemenman-Shafee-Bialek method (Nemenman et al., 2004) or some more recent methods using statistic copulas (Ince et al., 2017; Safaai et al., 2018). Each method has its advantages and downsides (see (Panzeri et al., 2007) for a comparison of some of them), which leaves the experimenter puzzled and in dire need of a mathematician (Borst and Theunissen, 1999; Magri et al., 2009; Piasini and Panzeri, 2019; Timme and Lapish, 2018).

However, there is another way to calculate the entropy of a signal, through what is called the Source Coding Theorem (Cover and Thomas, 2006; Larsson, 1996, 1999; Shannon, 1948; Wiegand, 2010). In signal processing, data compression is the process of encoding information using fewer bits than the original representation. In case of lossless compression, it does so by sorting parts of the signal by their redundancy and replacing them by shorter code words (Huffman, 1952; Shannon, 1948). The Source Coding Theorem specifies that a signal of size *S* and of entropy *H* cannot be compressed into less than *S*\**H* bits without losing information. Therefore, with a perfect lossless compression method the size of the compressed signal is proportional to the original signal entropy (Cover and Thomas, 2006; Larsson, 1996, 1999; Shannon, 1948; Wiegand, 2010). This method has been widely described in the field of physics, where estimating entropy via compression algorithms has been done several times (Avinery et al., 2019; Baronchelli et al., 2005; Martiniani et al., 2019, 2020) but to our knowledge it has been used only twice in the field of neurosciences (Amigó et al., 2004; London et al., 2002) to estimate the entropy of spike trains and the information efficacy of a synapse.

When choosing this way of calculating entropy, the choice of the compression algorithm becomes critical as the compressed signal must be the smallest possible in order to represent the entropy of the original signal. It is of course possible to craft its own compression algorithm (see (London et al., 2002)), but thankfully this application has been broadly used in the domain of informatics, in order to compress text and images efficiently on the hard drive of a computer or before sending data through a network. In particular, this led to the development of two principal entropy-coding compression algorithms: the Huffman coding algorithm (Huffman, 1952) and the Lempel–Ziv– Storer–Szymanski algorithm (Storer and Szymanski, 1982; Ziv and Lempel, 1977), both used to compress text and image files. This Lempel-Ziv algorithm and its variants are the main tools for the estimation of entropy from data compression (Avinery et al., 2019; Baronchelli et al., 2005; Benedetto et al., 2002; Martiniani et al., 2019).

Portable Network Graphics (or PNG, see specifications at https://www.w3.org/TR/PNG/ or http://www.libpng.org/pub/png/) is a graphic file format supporting lossless data compression. Its high versatility and fidelity made it widely used for saving and displaying pictures. Its lossless compression is based on the combination of the Lempel–Ziv–Storer–Szymanski and Huffman algorithms and is called DEFLATE (Deutsch, 1996). In short, it consists of two main steps: i) bit reduction, replacing commonly used symbols with shorter representations and less commonly used symbols with longer representations by Huffman coding and ii) Duplicate string elimination by detecting duplicates and replacing the occurrences by a reference to the first one, by LZSS algorithm. Its great efficacy made it a reference for comparison with other entropy-encoding image compression methods (Bian et al., 2019; Cover and Thomas, 2006; Hou et al., 2020; Mentzer et al., 2020) and it is even used directly to estimate image entropy (Wagstaff and Corsetti, 2010).

In this paper, we show that measurement of PNG file output size of neuroscientific data (in Bytes on the hard drive) is a reliable and unbiased proxy to estimate the level of entropy of electrophysiological or morphological data. First, the relationship between entropy level and file size is linear. Second, by simply dividing the size of the PNG file (in Bytes) by the number of pixels in the image, we can obtain a “PNG Size Rate” (or “PNG Rate”, in Bytes per pixels), which slowly converges towards a stable value when increasing the amount of recorded data. This allows the experimenter to compare multiple recordings of different sizes. Therefore, even if the PNG Rate does not provide the exact entropy value, it is robust enough to estimate the level of entropy in response to different experimental conditions. Furthermore, with minimal modifications of the PNG file, we validate estimation of the mutual information between a stimulation protocol and the resulting experimental recording.

## Methods

### Calculation of entropy rate and mutual information rate via the direct method and quadratic extrapolation

We calculated the entropy and mutual information rates using the direct method and quadratic extrapolation, as originally described by (Strong et al., 1998) and recently used by (Juusola and de Polavieja, 2003; Panzeri et al., 2007; de Polavieja et al., 2005). We take their definition of entropy rate *R*, as the average entropy per symbol. We use their terminology for naming the different parameters as well.

The number of possible combination symbols is defined by *T* in this manuscript, so we can write: 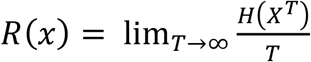. If *T* = 1, all the time points are considered independent, if *T* = 2, the time points are grouped by 2, if *T* = 3, the time points are grouped by 3, etc… Moreover, the entropy is depending on two more parameters: the portion of the signal considered (parameter *Size*: if *Size* = 1, the whole signal is considered, if *Size* = 0.5, half of the signal is considered, etc.) and the number of quantization levels (parameter *v*: if *v* = 2, the points are put in 2 classes of amplitude, if *v* = 10, the points are put in 10 classes of amplitude, etc.). As we need to estimate the entropy rate of our signal *R*_*S*_ for infinite possible quantization levels and infinite size of the signal, this yields:

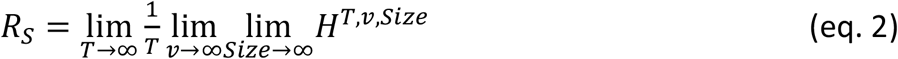

The direct method with quadratic extrapolation consists in two steps:

i. The first step of this method, comparable to a “brute force” approach, is to calculate the different values of entropy rates of the signal for multiple probability spaces. This means varying the portion of the signal, the quantization range and the possible combinations of values of the signal. As described in the introduction, each modification of *Size, v* or *T* will give a different entropy value. Ideally, we would like to calculate it for every possible value of *Size, v* and *T*, but this is not possible in practice: we want an estimation of entropy rate for recordings of infinite size, quantization range and combination bins. We thus limit ourselves to probable combinations for this first step, keeping in mind we will need a set of values big enough to yield accurate fits in the second step. For example, in figure 1, to estimate the entropy rate on different length of recordings the parameter *Size* was successively set as 1, 0.9, 0.8, 0.7, 0.6, 0.5, which takes decreasing portions of the signal, from full signal to half of it. As we are measuring the entropy rate from a discrete uniform distribution (white-noise-like) with 2 to 256 possible values, we successively set the quantization range *v* as 2, 4, 8, 16, 32, 64, 128, and 256. Any range higher than 256 will yield the same value of entropy rate. The number of combination bins *T* was successively set as 1, 2, 3, 4, 5, 6, 7, 8, in order to measure the entropy with no combinations (*T* = 1) until 8 possible combinations of values (*T* = 8). This produced 6 * 8 * 8 = 384 distinct values of entropy for every trial. Entropy values for different trials of the same condition were averaged together.
ii. The second step (Figure 1B) will extrapolate these data to find their limit value to infinite size, quantization range and combination bins. The previously calculated values are first plotted against 1/*Size* and the intersections to 0 estimated by quadratic fit of the data. This gives us the entropy values for every *v* and *T*, corrected for infinite *Size* of the recordings. These values are then quadratic fitted against 1/*v*. The intersection to 0 gives us entropy values for every number of combination *T*, corrected for infinite *Size* and infinite number of quantization levels *v*. As combinations of *T* elements will happen 1/*T* times, we divide these values by *T* and these new values are fitted against 1/*T* to estimate the extrapolation to 0. By performing this triple extrapolation and dividing by the number of combination bins, we can estimate the entropy rate *R*_*S*_ of the signal for theoretical infinite size of recording, infinite number of quantization levels and infinite number of combinations as in (eq. 2).

**Figure 1:**
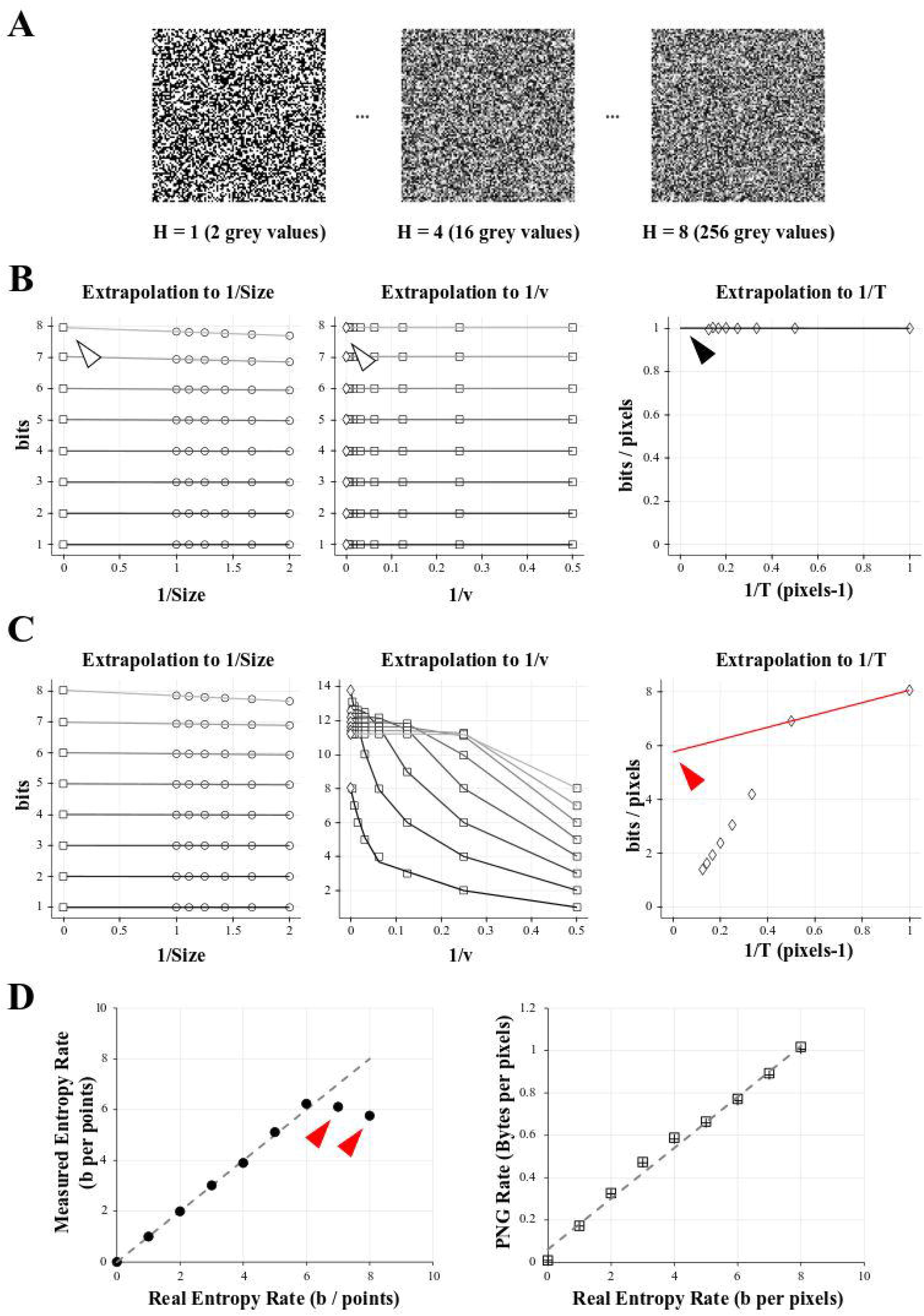
comparison of entropy and PNG file size on a model case. **A)** Examples of 10 000 data points of white-noise-like signal with growing number of grey levels and growing entropy (here showed as square picture signals). Left: 2 possible grey values, or entropy of 1 bit. Middle: 16 possible grey values, or entropy of 4 bits. Right: 256 possible grey values, or entropy of 8 bits. **B)** Direct calculation and quadratic extrapolations to 0 to calculate the entropy rate of the 1 bit-signal (left signal in A). Left: Plotting all the entropy values to 1/*Size* and extrapolating to 0 to get the value for infinite size (white arrowhead). For clarity, only the condition for *v* = 2 is shown (but this was done for *v* equal to 2, 4, 8, 16, 32, 64, 128, and 256). Middle: Plotting the limits values obtained for 1/*Size* = 0 (left graph) versus 1/*v* and extrapolating to 0 to get the value for infinite number of quantization levels (white arrowhead). Right: Plotting the limits values obtained for 1/*v* =0 (middle graph) versus 1/*T* and extrapolating to 0 to get the value for infinite number of combinations bins (black arrowhead). Note that this value is close to 1 bit/pixel, as expected when using a signal made of uniform white noise with 2 possible values. **C)** Direct calculation and quadratic extrapolations to 0 to calculate the entropy rate of the 8 bits-signal (right signal in A). Left: Plotting all the entropy values to 1/*Size* and extrapolating to 0 to get the value for infinite size. For clarity, only the condition for *v* = 2 is shown (but this was done for *v* equal to 2, 4, 8, 16, 32, 64, 128, and 256). Middle: Plotting the limits values obtained for 1/*Size* =0 (left graph) versus 1/*v* and extrapolating to 0 to get the value for infinite number of quantization levels. Right: Plotting the limits values obtained for 1/*v* = 0 (middle graph) versus 1/*T* and extrapolating to 0 to get the value for infinite number of combinations bins. Note that in the final graph (right), points do not follow a linear trend. When using the last 2 points for extrapolation to 0, we obtain a value of 5.6 bits/pixels (red arrowhead), far from the expected value of 8 bits. This reveals the sampling disaster (not enough points in the signal to properly estimate the entropy) **D)** Left: When plotted against the real entropy value, the direct method with quadratic extrapolation shows examples of sampling disaster for high values of entropy (red arrowheads). Right: When simply saving all the signals described in A) and dividing their size by the number of pixels in the image we obtained the PNG Rate in Bytes per pixel. The PNG Rate shows a linear relationship with the Entropy Rate (*y* = 0.12 *x* + 0.06, R^2^ = 0.99). Note that this is true for pictures made either of square signals (squares plot) or linearized signals (crosses plot).

To obtain *R*_*N*_ in Figure 4, the entropy rate of the noise, instead of calculating the entropy H along the length of the signal, we did it at every time point across the successive trials. This is equivalent to simply transpose the signal and re-applying the same method as for *R*_*S*_. Finally, we obtained the mutual information rate by subtracting *R*_*N*_ to *R*_*S*_ as

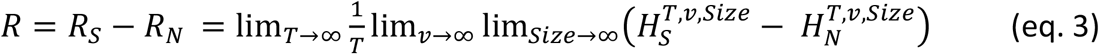

For Figure 4A, simulated recordings were down-sampled to 10 kHz before calculation of information rate. For Figure 4C, middle, simulated recordings were down-sampled to 3 kHz and binned as 0 & 1 depending of the presence of Action Potentials or not, similar to (London et al., 2002).

For figures 1B, 1C, 1D, 2, 3, 4A and 4B, The parameter *Size* was successively set as 1, 0.9, 0.8, 0.7, 0.6, 0.5; *v* successively set as 2, 4, 8, 16, 32, 64, 128, 256 and the parameter *T* was successively set as 1, 2, 3, 4, 5, 6, 7, 8. For figure 4C, the parameter *Size* was successively set as 1, 0.9, 0.8, 0.7, 0.6 and 0.5; *v* set as 2 and the parameter *T* was successively set as 1, 2, 5, 10, 20, 30 and 40. *R*_*S*_, *R*_*N*_ and information transfer rate were calculated by direct method and successive quadratic extrapolations, as described above.

All of this was done by custom scripts written in Python 3.7 Anaconda with Numpy, Pandas and pyABF modules. These scripts are available in Python and Labview format at https://github.com/Sylvain-Deposit/PNG-Entropy.

### Export to PNG format

Export to PNG was made with 3 different softwares: i) Anaconda 3.7 (https://www.anaconda.com/) and the pyPNG package (https://pypi.org/project/pypng/) for Figures 1, 2A, 3A, 3B ; ii) Labview 2017 and Vision 2017 (National Instruments) for Figures 2B, 3C, 4A-C, 5C; iii) the FIJI distribution of ImageJ software (Rueden et al., 2017; Schindelin et al., 2012) for figure 6C. Signals were normalized to 256 values from 0 to 255 simply by subtracting the minimal value of the signal, then dividing by the maximal value and multiplying by 255. It was then saved as PNG format in 8-bits range (256 grey values). For figure 4C, as the signal was binarized we saved it with a 1-bit range (2 grey values). This script and others are available in Python and Labview format in a GitHub depository: https://github.com/Sylvain-Deposit/PNG-Entropy

**Figure 2:**
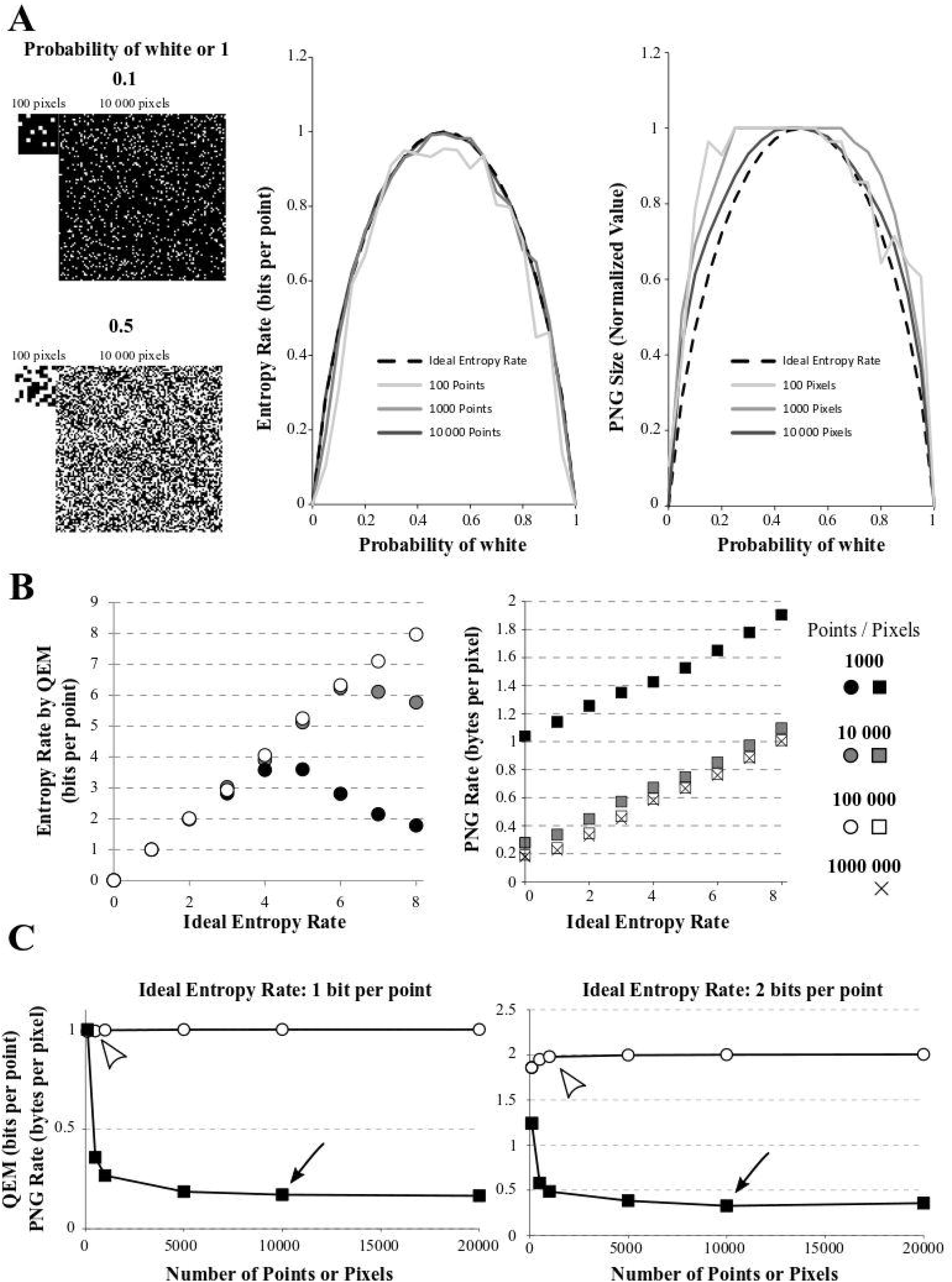
Effect of signal size on quadratic extrapolations method and PNG Rate. **A)** Left: The signals have 2 possible values (1 or 0, here shown by black or white) and can be of length 100, 1000 or 10 000 data points (only 100 and 10 000 points signals are shown). The signals with growing amount of white points (from 0% to 100%). Here, example of 10 % of white points (up) or 50% of white points (down) are shown. Middle: In dashed line: the ideal bell-shaped curve calculated by the Shannon’s formula. When using the quadratic extrapolations method, we needed at least 10 000 points to converge to the ideal bell-shaped curve (compare dark grey line to black dashed line). However, if our signal is short, it is impossible to calculate high entropy values (100 points for example, light grey curve). Right: normalized PNG file sizes of the same signals, showing that when we increase the number of points in the signal, we progressively fit the obtained curve toward the ideal entropy curve. This convergence is slower than when using the quadratic extrapolations method. **B)** Effect of the number of points on entropy calculation for white-noise signals. The different signals differ by their number of possible values (from 1 to 256 possible values leading to ideal entropy rates from 0 bits/point to 8 bits/point) and by their number of points (from 1000 to 1 000 000 points). Left: Effect of the number of points on the entropy rate calculation via the quadratic extrapolations method (QEM). For signal with few points (1000 and 10 000 points), the curve shows a biphasic behavior revealing the sampling disaster: a signal of few points cannot be used to calculate high entropy values. When increasing the number of points, the curve becomes accurate for every entropy value. Right: same demonstration when using the PNG Rates. Even with few points, the curve shows a linear behavior. However, increasing the number of points modifies the slope and intercept of the linear curve, until it stabilizes to an optimal solution. **C)** Effect of the number of points on entropy calculation for white-noise signals. The different signals differ by their number of possible values (2 or 4 possible values leading to ideal entropy rate from 1 bit/point or 2 bits/point) and by their number of points (from 100 to 20 000 points). Left: Calculations for the 1 bit/point signal. Using the quadratic extrapolations method (QEM, white circles), the calculated entropy rate versus the number of points in the signal shows a quick convergence to the optimal value (white arrowhead, around 500 points, value of 0.9923 bits per point). If we plot the PNG Rate versus the number of points in the signal (black squares), we obtain a curve decreasing slowly to a stable value (Black arrowhead, around 10 000 points, value of 0.17 Bytes per pixels). Right: Same calculations made with a signal of 2-bits/point signal (4 different values possible). The direct method converges in around 1000 points (calculated value: 1.9794 bits per points) while the PNG Rate converges in around 10 000 pixels (calculated value: 0.32 bytes per pixels).

A minimal file of PNG format is composed of a header and several parts of data, named critical chunks (https://www.w3.org/TR/PNG/#5DataRep). To these minimum requirements it is possible to add ancillary chunks (https://www.w3.org/TR/PNG/#11Ancillary-chunks) containing various information such as Software name, ICC profile, pixels dimensions, etc… If useful, this is hindering the estimation of entropy as it represents an overhead to the final size of the file. To estimate this overhead for each of our software we saved an image of 100 * 100 values of zeros, which corresponds to black in 8-bits grey levels and has an entropy of 0. With pyPNG, Fiji and Labview we obtained three PNG files of size 90, 90 and 870 Bytes, respectively. When repeating the experiment of figure 1, we obtained similar linear fits of slopes 1.21 (R^2^ = 0.99), 1.18 (R^2^ = 0.99) and 1.21 (R^2^ = 0.99) respectively.

For Figure 6C, we used Fiji for every image of the collection and we: i) extracted the channel number 2 containing the MAP2 staining; ii) converted the file to 8-bits grey levels; iii) thresholded it to remove every intensity values under 10 to remove most of the background; iv) saved the new file as PNG format, v) checked the size of this new file and vi) divided the size in kBytes by the number of soma visible in the field.

### Neuronal modeling

A single compartment model was simulated with NEURON 7.7 (https://www.neuron.yale.edu/neuron/). All simulations were run with 100-μs time steps. The nominal temperature was 37°C. The voltage dependence of activation and inactivation of Hodgkin-Huxley–based conductance models were taken from (Hu et al., 2009) for g_Nav_ and g_KDR_. The equilibrium potentials for Na^+^, K^+^, and passive channels were set to +60, −90 and −77 mV, respectively. The conductances densities were set to 0.04 S/cm^2^, 0.01 S/cm^2^ and 3.33*10^−5^ S/cm^2^ for g_Nav_ and g_KDR_ and passive channels, respectively.

The model was stimulated using various numbers of excitatory synapses using the AlphaSynapse PointProcess of the NEURON software. The time constant and reversal potential were the same for every synapses and were set to 0.5 ms and 0 mV, respectively. The size of EPSPs produced by the synapses were randomly chosen using a lognormal distribution of EPSPs amplitude experimentally described in L5 pyramidal neurons (Lefort et al., 2009). Each synapse stimulated the model once during a simulation and the time onset was randomly chosen.

For the simulations of Figure 3, the number of synapses simulating the model depended on the duration of the simulation and spiking frequency desired. For example, to calculate the entropy rate in the case of a 1 Hz spiking frequency during 400 seconds, we simulated the model with 3200 of the synapses described above. For higher spiking frequencies, the number of synapses was increased.

**Figure 3:**
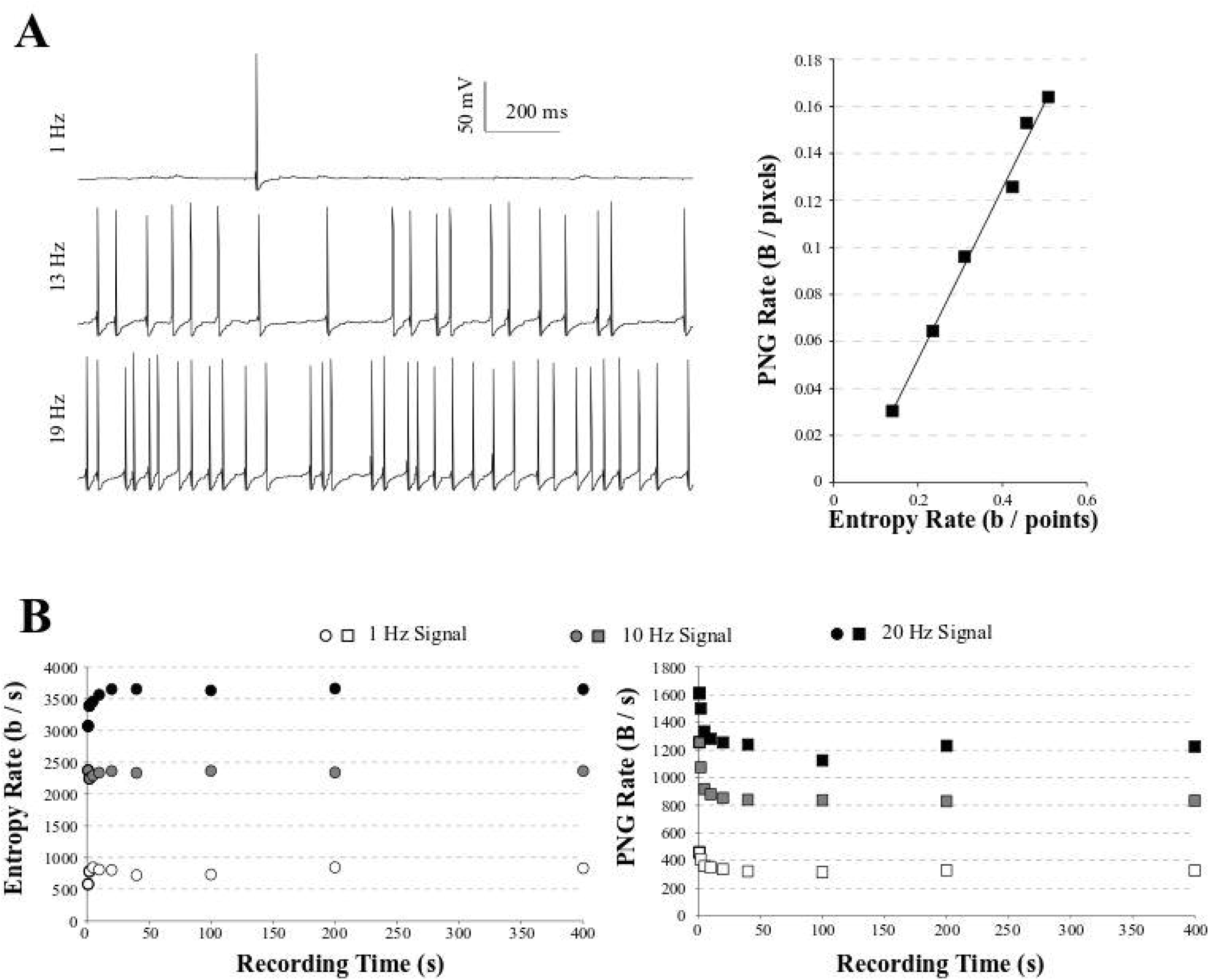
PNG Rate on a model of neuronal activity. **A)** Left: example of traces obtained with our model, increasing the number of synapses to obtain spiking activity from 1 Hz (top) to 20 Hz (bottom). Right: The PNG Rate (Bytes/pixels) is linear to Entropy Rate (bits/points), following *y* = 0.36*x* − 0.06 (R^2^ = 0.99). **B)** Impact of the simulation duration on the Entropy Rate (Left) and PNG Rate (Right). Entropy Rate converges toward a stable solution in around 5 seconds of recordings, or 1 sweep. The PNG Rate needs around 40 seconds to obtain a stable value. As the PNG algorithm starts by linearizing the data, this is equivalent to 8 sweeps of 5 seconds each.

For the simulations of Figures 4A and 4B, each trace lasted 5 seconds. The number of synapses simulating the model depended on the spiking frequency desired. For example, to calculate the information transfer rate in the case of a 1 Hz spike train, we simulated the model with 400 of the synapses described above. We ran 20 trials of the simulation with the same train of synapses (Figure 4A). In order to introduce some jitter in the spiking times, we also injected a small gaussian current with a mean of 0 nA and a standard deviation of 0.0005 nA during the 5 seconds of the simulation. We reproduced this whole protocol for others desired spiking frequency, using increasing number of synapses (for example: 1900 synapses for a 19 Hz spiking).

**Figure 4:**
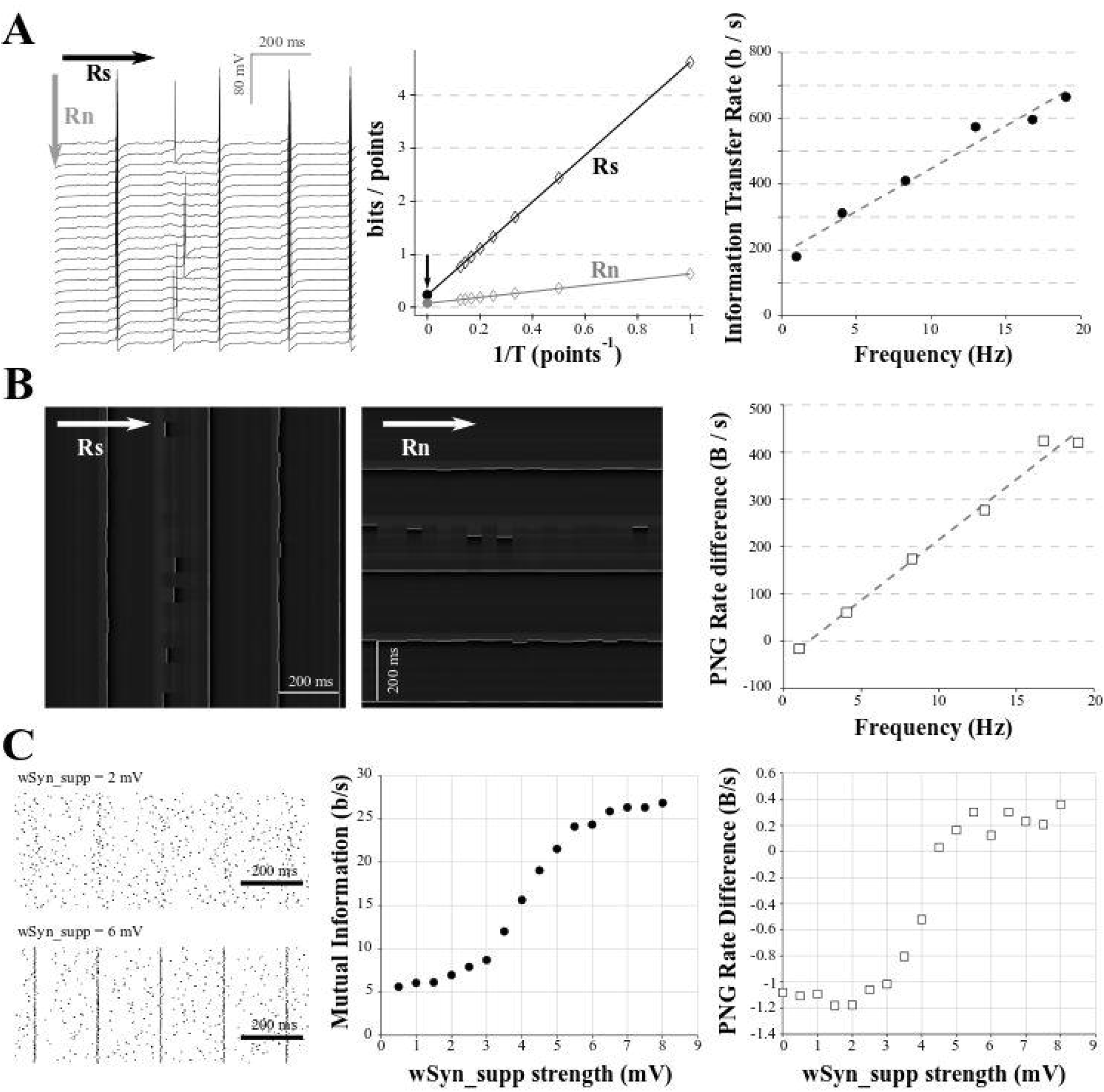
Comparison of Mutual Information and PNG Rate on a neuronal model. **A)** We considered different amount of synapses to obtain different spiking frequencies (from 1 to 20 Hz). For each spiking frequency, the model was run for 20 trials with the same synaptic inputs. Due to the injection of a small Gaussian noise current, we obtain variability in the spiking of the different trials. Left: example of 20 generated trials with a 5 Hz spiking activity. Note that despite the same synaptic activity between trials, the small Gaussian noise current induces variability in the spiking of the different trials. Arrows show the direction used with the quadratic extrapolations method to calculate the signal entropy *R*_*S*_ and noise entropy *R*_*N*_. Middle: calculation of the Entropy Rate of the Signal *R*_*S*_ and the Entropy Rate of the noise *Rn* for the full voltage of the cell for each condition. The Information transfer Rate *R* is the difference between *R*_*S*_ and *R*_*N*_. Right: Information transfer Rate (*R*_*S*_-*R*_*N*_) between the synaptic stimulation and the neuronal activity, plotted versus the spiking frequency. This follows a linear trend (*y* = 25.935 *x* + 188.15, R^2^ = 0.97). **B)** Left: Conversion of the modeled traces in A) as a 256 grey values PNG file. As the PNG conversion algorithm is line-wise, we have to save the image a first time, divide by the number of pixels and multiply by the sampling rate to get the PNG Rate of the signal in Bytes/s (equivalent to Rs in the quadratic extrapolation method) (Left). Then we have to rotate the image 90 degrees, save it a second time, divide by the number of pixels and multiply by the sampling rate to get the PNG Rate of the noise in Bytes/s (equivalent to Rn in the quadratic extrapolation method) (Middle). Arrows show the direction of compression. Right: the subtraction of the PNG Rate of the signal and the PNG Rate of the noise follows a linear trend with the spiking frequency (*y* = 25.55 *x* − 41.2, R^2^ = 0.98). **C)** The model was run with the same amount of background synapses to obtain a spiking frequency of 5 Hz. The onset of the background synapses was chosen randomly at each trial to create variability in the spiking from trial to trial. A supplementary synapse was added and stimulated the model every 200 ms. The strength of the supplementary synapse varied from 0.5 to 8 mV. 100 trials were run for each strength of the supplementary synapse. Left: examples of rastergrams showing the impact of the supplementary synapse on our neuronal model. With a low synaptic strength (*wSyn_supp* = 2 mV, top), this synapse barely drives the model spiking. With a high synaptic strength (*wSyn_supp* = 6 mV, bottom), the neuron spiking is synchronized with the occurrence of the synapse. Middle: Information transfer Rate between the strength of the supplementary synapse and the neuronal spiking. As expected, it follows a sigmoidal behavior. Right: rastergrams were saved as PNG files, divided by the number of pixels and multiplied by the sampling rate to obtain the PNG Rate of the signal. Then, the rastergrams were rotated 90 degrees, saved again, divided by the number of pixels and multiplied by the sampling rate to obtain the PNG Rate of the noise. The difference of the 2 PNG Rates follows a similar curve as the Information transfer Rate.

For the simulations of Figure 4C, we stimulated the model with 750 of the synapses described above to get a spiking frequency around 5Hz. The time onsets and the amplitude of the synapses were randomly chosen at each simulation. We also added one supplementary synapse (Syn_supp) which stimulates the model every 200 ms (i.e 25 times in 5 s). The size of the EPSP size produced by this synapse was called *wSyn_supp*. When *wSyn_supp* was weak, this synapse did not drive the spiking of the model (Figure 4C, up left). When *wSyn_supp* was strong, this synapse drove the spiking of the model (Figure 4C, down left). We ran 100 simulations for each *wSyn_supp*.

### Place fields simulation

To simulate place fields recordings, we programmed a simple random-walk in a 20*20 cells area with a place field at its center. A mock animal was placed randomly in this area and allowed to “run” for a finite set of step (typically, 10 000 steps). For each step of the simulation, the animal could choose uniformly between one of the 8 neighboring cell or stay in the same cell. For each step of the simulation, the spike frequency of a recorded place field was generated by calculating the distance *x* of the animal from the place field center and indexing this distance to a sigmoid curve 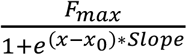 with *x*_*0*_= 7, *Slope* = 2 and *F _max_* = 10 to obtain a place field of roughly 3 cells of radius in the center of the simulated area (Figure 5). This script and others are available in Labview format in a GitHub depository: https://github.com/Sylvain-Deposit/PNG-Entropy

**Figure 5:**
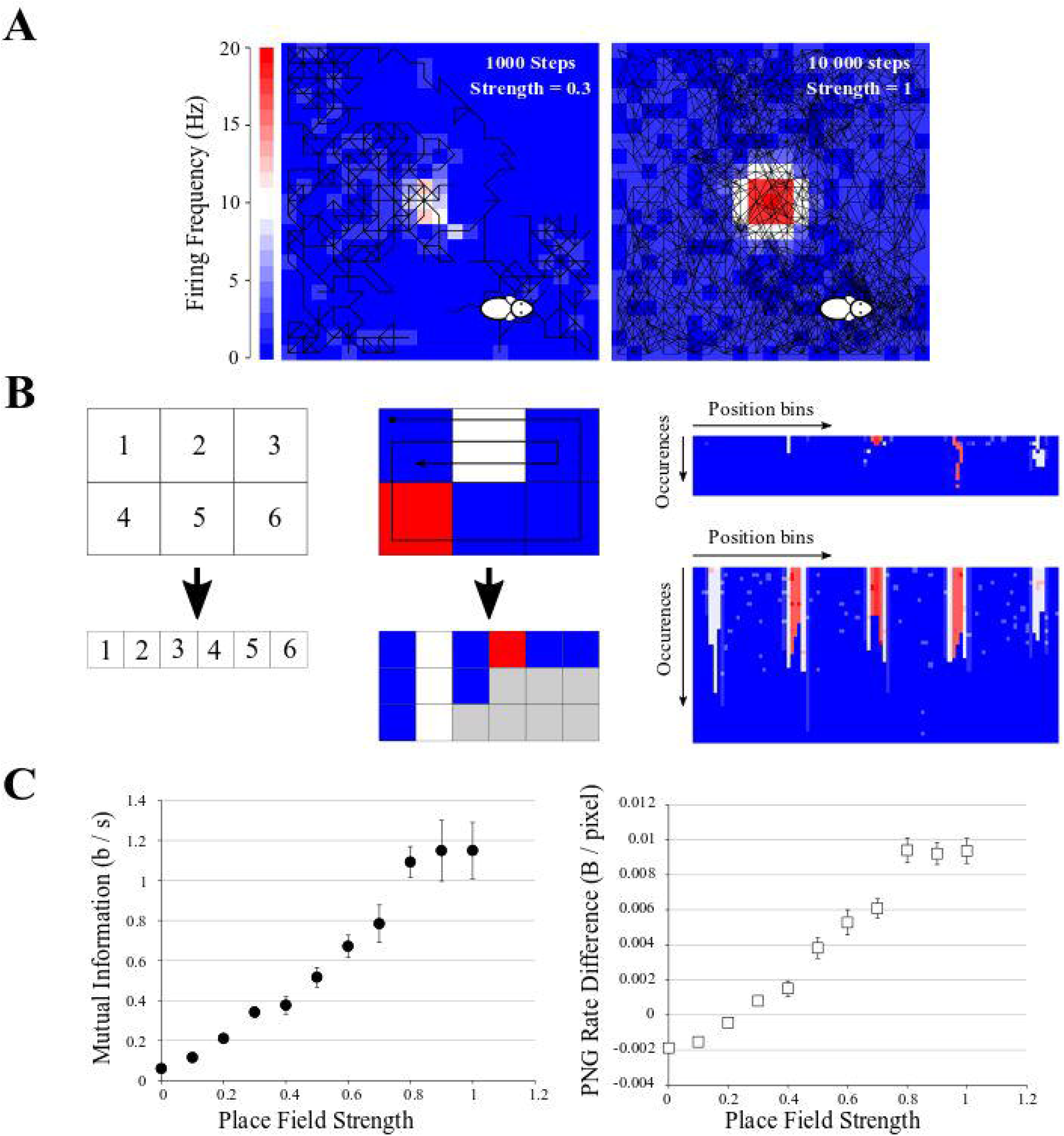
Application to Mutual Information from Place Fields. **A)** Examples of simulated data: a mock animal with a random walk in an enclosed environment. The firing rate of each space bin was modeled as a Gaussian-like centered in the field, with growing probability of firing. Left: example of a 1000-steps random walk with a place field of low probability of firing (*p* = 0.3). Middle: example of a 10 000-steps random walk with a place field of high probability of firing (*p* = 1). **B)** Simple explanation of the pairing function used to transform the Place field information to a 2D array to be saved in PNG format. Left, top: example of an area divided in 6 space bins, and each bin is given a unique index, which will be the index of the columns of the future image (left, bottom). Middle: if we follow the patch showed with the arrow, we traverse the indexes 1, 2, 3, 6, 5, 4, 1, 2, 3, 2, 1. For each traversed index, we place a pixel with the color corresponding to the firing frequency at the corresponding column index. If we go to the same position multiple times, we simply add multiple rows to the 2D array. There will be “holes” (grey squares) as the path did not go to every position the same number of times, but we can fill them with 0 (blue squares). We end up with a 2D array with the number of columns corresponding to the number of space bins (6 in this example) and the number of rows corresponding to the number of times the path entered the same space bin (3 for this short example, as the path entered the space bins 1 & 2 three times). Right: result of this pairing function on the 20*20 arena for the example seen in A), left (top) and A), right (bottom). These files were saved as PNG, rotated, saved again to obtain the difference in PNG Rates. **C)** Left: Mutual Information as *I*_*sec*_ (eq. 9) versus Place Field strength, showing a clear increase when the place field has a high chance to fire. Right: Difference in PNG Rates versus Place Field strength. Both curves are similar.

## Results

### Entropy estimation of white-noise-like signals

To test the usability of the PNG format to represent entropy, we started by generating 10 000 pixels of signal with a discrete uniform distribution (white-noise-like) which could take 2 values (Figure 1A, 2 grey values). As we are using a uniform distribution, we fully know the probability distribution (in that case, *p*(*xi*)=1/*N*), we can apply the eq. 1 and obtain an entropy *H* of 1 bit. We then repeated this noise model progressively increasing the number of possible values by power of 2 until 256 (i.e 4 possible values, 8 possible values, 16 possible values etc… up to 256 possible values). The 256 possible values of uniform distribution correspond to an entropy of 8 bits (Figure 1A). The entropy rate *R* is defined 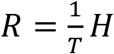 as with *T* being the sampling of the signal. In this model case, we can take *T* = 1 (i.e a sampling of 1 pixel) to finally obtain an entropy rate in bits per pixel.

As a control way to calculate the entropy of our signals, we used a method described in (Juusola and de Polavieja, 2003; Panzeri et al., 2007; de Polavieja et al., 2005; Strong et al., 1998), involving the use of the Shannon’s formula in many probability spaces followed by quadratic extrapolations to estimate the entropy rate in a case of an infinite size signal, an infinite quantization and an infinite time sampling, see Methods for explanations. This method allows the exact calculation of the signal entropy rate but is long to compute and needs programming skills to be implemented. It will be used all along this study as a control to be compared with the PNG method. It will be referred as “the quadratic extrapolations method”.

In our simple model data, we can see that when using only 2 different values for the white-noise-like signal (Figure 1B), the quadratic extrapolations method gives an entropy rate close to 1 bit/pixels as expected (Figure 1B, right panel, black arrowhead).

However, when increasing the number of possible values until 256 (real entropy rate *H* of 8 bits/pixels) we start to unravel the so-called “sampling disaster” (Figure 1C). In fact, the quadratic extrapolations method gives a value around 5.7 bits/pixels (Figure 1C, right panel, red arrowhead), far from the expected one. This is easily explained by the number of data points we have in our model signal. The last extrapolation concerns the *T* combinations of values to calculate the entropy rate and when using a uniform distribution and with *T* equal to 1, it means we need at least 2^8^ = 256 points to properly estimate the probability distribution. However, if *T* increases to 2 (1/*T* = 0.5 in the Figure 1C), we then need at least 2^8*2^ = 65536 points to estimate the probability distribution, when we had only 10000. The probability distribution is thus insufficient to properly estimate the entropy rate of this signal with 256 values. Even by using quadratic extrapolations to compensate for the sampling bias, we can see it gives a wrong result for high entropy values when there are not enough data points (red arrowheads in Figure 1D, left panel).

As a comparison, we simply saved the 10 000 pixels data of increasing white-noise-like signal (Figure 1A) as PNG image file with a depth of 8 bits (and thus 256 possible values). We measured the space taken by these files on the hard drive (in Bytes) and divided this value by the number of pixels to obtain the PNG Rate (in Bytes/pixels) (Figure 1D, right panel). The PNG Rate was linearly correlated with the real entropy rate (*y* = 0.12 *x* + 0.06, R^2^ = 0.99). The PNG compression algorithm works with linearized data, which means there is no difference when saving pictures as a 100 * 100 square format (figure 1D, right panel, squares plot) or saving a 10 000-single line (same panel, crosses plot). From this first test, we can conclude that for our model signal, the size of a PNG file divided by the number of pixels has a linear relationship with its entropy rate.

To better investigate the effect of recording length on entropy rate and PNG conversion, we generated 100 points of white-noise-like signal with 2 possible values (0 and 1) and progressively increased the percentage of 1 in this signal, from 0 to 100% (0 is represented by a black pixel, 1 is represented by a white pixel) (Figure 2A). As we know the ideal probability distribution of this signal, we can use the Shannon’s formula (eq.1) to calculate the real entropy rate and obtain the bell-shaped curve as described by (Shannon, 1948) (Figure 2A, middle, black dashed curve). However, we see that if we calculate the entropy rate from our 100 points signal with the quadratic extrapolations method (Figure 2A, middle, light grey), the curve obtained is far from ideal: it is unable to reach the proper value of 1 for maximum entropy. This is another example of sampling disaster. We have to increase the number of points in our signal up to 10 000 points to obtain a maximum entropy rate corresponding to the maximum ideal entropy rate (Figure 2A, middle, middle grey and dark grey). Then, we saved the same signals to PNG files, measured their size on the hard drive and divided by the number of pixels to obtain the PNG Rates of each signal. Finally, we normalized the obtained curves to compare their shapes (Figure 2A, right). We can see that the obtained curve progressively fits the ideal entropy rate curve when the number of points increases (Figure 2A, right), but at a slower rate than the direct method. In fact, even with 10 000 points, the curve obtained by the PNG method does not fit perfectly the ideal curve.

As an illustration of the effect of the number of points on each method, we generated white-noise-like signals with increasing number of possible values from 0 to 256 (thus entropy rates from 0 to 8 bits per point) and we increased the number of points in these signals from 1000 to 1000 000 points (Figure 2B). We used the quadratic extrapolations method to calculate the entropy rates, and saved the same signals as PNG files to obtain the PNG rate. When plotted against the ideal entropy rate, we can see that these two methods yield different behaviors (Figure 2B). When calculating the entropy rate by the quadratic extrapolations method, the sampling bias dramatically modifies the high values of entropy rate and we obtain biphasic curves (Figure 2B, left). This indicates that we cannot estimate high entropy values with short signals. However, the values are perfectly accurate for low entropy rates or if the number of points in the signal is sufficient (around 100 000 points in this case). When plotting the PNG rate, we can see that the relationship between PNG Rate and the ideal Entropy Rate is always linear, but the number of points affects the slope and intercept of the obtained curve (Figure 2B, right). This relationship converges towards a stable curve for long signals (around 100 000 pixels). This means that if we have multiple signals of the same size, PNG Rate can properly estimate the modifications of entropy rate between them as the relationship PNG Rate/entropy rate is linear. However, if we have signals of different sizes, we have to check before if the PNG rate converged to a stable value to be able to compare them.

To estimate the number of points needed for a correct entropy rate estimation of a given signal, we generated white-noise-like signals of 100, 500, 1000, 5000, 10 000 and 20 000 points. Those signals could present either 2 possible values (1 bit per point, Figure 2C, left) or 4 possible values (2 bits per point, Figure 2C, right). First, we calculated their entropy rates by the quadratic extrapolations method and plotted them versus their number of points (Figure 2C, white circles). When using a white-noise-like signal with 2 possible values (ideal entropy rate of 1 bit per point), 500 points were enough to find the optimal value of entropy rate by the quadratic extrapolations method (Figure 2C, left, white arrowhead). If we increase the number of values in our generated signal to 4 possible values (ideal entropy rate of 2 bits per point), the quadratic extrapolations method needs 1000 points to find the optimal value of entropy rate (Figure 2C, right, white arrowhead). As expected, when the entropy rate of the signal increases, the quadratic extrapolations method needs more points to find a stable value of the entropy rate. Second, we saved all the generated signals to PNG files, measured their size in Bytes on the hard drive and divided these sizes by the number of pixels in the image file to get the PNG Rates of the signals. Then, we plotted the PNG Rates versus the number of points in the signals (Figure 2C, black squares). When using a white-noise-like signal with 2 possible values (ideal entropy rate of 1 bit per point), we can see that the PNG Rate reaches a stable value around 10 000 pixels (Figure 2C, Left, black arrow). With a white-noise-like of 4 possible values (ideal entropy rate of 2 bits per point), the PNG rate reaches a stable value around 10 000 pixels as well (Figure 2C, Right, black arrow). By this experiment, we concluded that both methods converge towards a stable solution when the number of points in a signal increases. However, the quadratic extrapolations method is much quicker to converge than the PNG Rate method.

From these examples, we conclude that the number of points in the signal is critical to capture its entropy rate. In the case of short signals, the quadratic extrapolations method gives biphasic relationships with the Entropy Rate, making it unsuitable to calculate high entropy values. The PNG Rate method stays linear to the Entropy Rate, which allows comparing files of the same size in a wide range of entropy values. Both methods converge to a stable solution when increasing the number of points in the signal, but the PNG Rate method convergence is slower. Both methods allow comparing multiple signals of different sizes, as long as the stable value is reached.

### Entropy estimation in electrophysiological signals

In order to test the ability of the PNG Rate method to estimate the entropy of electrophysiological signals, we created a single compartment model in NEURON 7.7. The model contained sodium and potassium voltage-dependent channels to allow the spiking (see Methods for model description). The model was stimulated using various numbers of EPSPs with amplitudes chosen randomly in a log-normal distribution described in (Lefort et al., 2009). By increasing the number of synapses stimulating the model, we could increase the spiking frequency.

We ran simulations of 5 seconds with an increasing number of synapses, to obtain model traces ranging from 1 to 20 Hz AP frequency (Figure 3A, left). For each spiking frequency condition, we ran 20 trials with the same synaptic inputs (a small gaussian noise current was added to induce variability among trials). We used the quadratic extrapolations method to calculate the Entropy Rate on all our model traces. In parallel, we saved these traces as 8-bits PNG files (256 possible values) and divided by the number of points in the image to obtain the PNG Rate. As expected, the PNG Rate has a linear behavior versus the Entropy Rate (Figure 3A, right). Therefore, the PNG Rate is suitable to estimate entropy rates variations in electrophysiological signals. As a control, we measured the Entropy Rate and the PNG Rate for different sizes of model traces at 1, 10 and 20 Hz, ranging from 1 second to 400 seconds. In our conditions, the Entropy Rate measured by the quadratic extrapolations method converges to a stable value in around 5 seconds of recordings (or 50 000 points, Figure 3B, left), whereas the PNG Rate needs around 40 seconds of recording (or 400 000 pixels, Figure 3B, right). These can be considered as the minimal time needed if one wants to compare multiple files of different sizes. Interestingly, as the PNG algorithm works with linearized data, there is no difference between 40 seconds of recordings and 8 sweeps of 5 seconds each. This means that in our model case, 20 trials of 5 seconds each were largely enough to estimate the stable value of the PNG Rate.

### Mutual Information estimation in electrophysiological data

Most of the time, the experimenter is not interested in the entropy itself, but in the mutual information between two variables *X* and *Y*. The mutual information *I*_*XY*_ measures the statistical dependence by the distance to the independent situation (Cover and Thomas, 2006; Shannon, 1948) given by

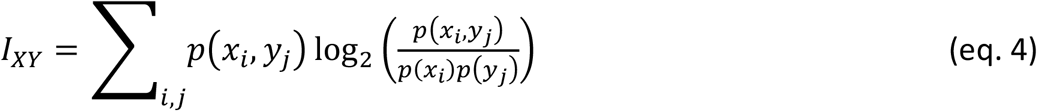

Therefore, when *X* and *Y* are independent *p*(*x*_*i*,_ *y*_*j*_) = *p*(*x*_*i*_)* *p*(*y*_*j*_), so *I* _*XY*_ = 0 bits.

The mutual information can also be rewritten as the difference between the entropy of *X* and the conditional entropy of *X* given *Y*:

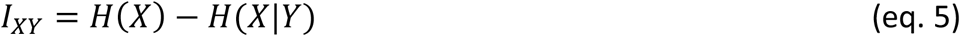

Where *H(X)* is the entropy already described (eq.1) and

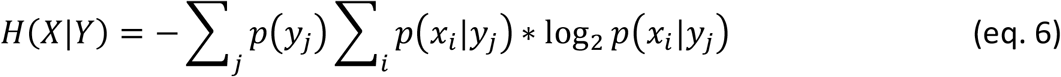

If a neuron is stimulated several times by the stimulus, we can define *X* as the response of the neuron to the stimulus and *Y* as the stimulus received by the neuron (Borst and Theunissen, 1999). In that case, *H(X)* is the entropy of the neuronal response (quantifying the total variability of the neuronal response) also called the total entropy: *H*_*S*_. *H(X*|*Y)* is the entropy of the neuronal response given the stimulus. As the same for each trial, the only source of variability is the intrinsic noise of the neuron. In that case, *H(X*|*Y)* can be interpreted as the noise entropy H_N,_ quantifying the variability of the neuronal response which is still present even if the neuron is submitted to the same stimulus at each trial. So, in the case of a neuron stimulated with the same stimulus during different trials, *H*_*S*_ quantifies the average variability of the neuronal response during one trial and *H*_*N*_ quantifies the variability of the neuronal response across the trials. In that case the mutual information between the stimulus and the neuronal response is given by:

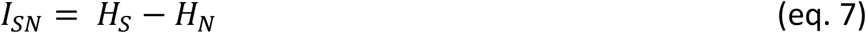

Where *Hs* is the averaged entropy of the neuronal response over every trial and:

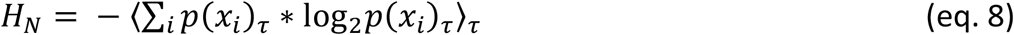

where *p*(*x*_*i*_)_τ_ is the probability of finding the configuration *x*_*i*_ at a time ⍰ over all the acquired trials of an experiment (Juusola and de Polavieja, 2003; de Polavieja et al., 2005; Strong et al., 1998). Finally, we can obtain the information transfer rate *R*, by using the quadratic extrapolations method and dividing by the time sampling (Juusola and de Polavieja, 2003; de Polavieja et al., 2005; Strong et al., 1998), as:

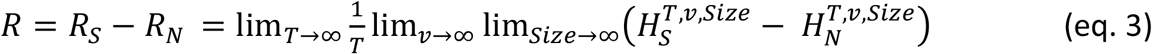

where *R*_*S*_ is the entropy rate of the signal and *R*_*N*_ is the entropy rate of the noise (see Methods for description of the quadratic extrapolations method). In practical terms, this means to acquire multiple recordings of the same experiments, apply the quadratic extrapolations method first on each trial and average the results to obtain *R*_*S*._ Then, to apply the same method across the trials for every time point ⍰⍰ and average the results to obtain *R*_*N*_. The information transfer rate *R* is thus the rate of the mutual information transfer between the stimulation protocol and the acquired trials.

In order to apply this method to electrophysiological signals, we used the same model as in the paragraph 3.2. In a first step, the synapses number was chosen to obtain a spiking frequency of 1 Hz. We ran 20 trials of 5 seconds with the same train of synapses (Figure 4A, left). In order to introduce some randomness in the spiking between trials, a small Gaussian noise current was also injected (Figure 4A, left). We calculated *R*_*S*_ and *R*_*N*_ using the quadratic extrapolations method. (Figure 4A, middle) and subtracted *R*_*N*_ to *R*_*S*_ to obtain the information transfer rate (here presented in bits/s) (Figure 4A, right). We then reproduced this protocol with various numbers of synapses to obtain different spiking frequencies. As expected, when we increased the number of synapses, we increased the spiking frequency and the information transfer rate between our stimulation and the response (Figure 4A, right). This measure follows a linear trend, similar to previous results obtained in literature (Juusola and de Polavieja, 2003; de Polavieja et al., 2005).

As already described (Figure 1D, right), the PNG format is line-wise. The compression algorithm will thus be sensitive to the orientation of the image we have to compress. To estimate the PNG Rate of the signal, we converted our voltage signals to an 8-bits PNG image (256 levels of grey). As our signals are 20 trials of 5s at 10 kHz sampling, this yielded a 1 000 000 pixels of 256 grey scale image (Figure 4B, left). We saved this first version of the image, measured the size of the files on the hard drive, divided this number by the number of pixels in the image and multiplied the value by the sampling (10 kHz) to obtain the PNG Rate of the signal in Bytes/s. To estimate the PNG Rate of the noise, we simply rotated the image 90 degrees and saved it again to PNG format. This rotation constrains the algorithm to calculate the entropy through the acquired trials and not through the signal itself, thus estimating the entropy rate at each time point across all the trials (as for *R*_*N*_ in the quadratic extrapolations method) (Figure 4B, middle). We measured the size of the newly generated file, divided again by the number of pixels in the image and multiplied the value by the sampling (10 kHz) to obtain the PNG Rate of the noise in Bytes/s. Finally, we subtracted the PNG Rate of the noise to the PNG Rate of the signal. As we can see, this difference of PNG Rates follows a linear behavior, increasing with AP frequency similarly to the direct measure of the information transfer rate (Figure 4B, right). We concluded that the PNG method allows accurate estimation of information transfer rate modifications through different conditions in electrophysiological data (here different spiking frequencies).

### Synaptic information efficacy estimation

As a second example, we reproduced the protocol made by (London et al., 2002) to estimate the information transfer between one synapse and the postsynaptic neuron spiking (also called Synaptic Information Efficacy, SIE). In this study, the authors showed that a larger synapse drove the postsynaptic spiking in a greater manner, which increases the SIE. To reproduce this result, we used the model described in the previous paragraphs. We stimulated the model during 5 seconds with 750 synapses to get a spiking frequency around 5Hz. Moreover, we added a supplementary synapse stimulating the model regularly every 200ms. When the EPSP size of this supplementary synapse (*wSyn_supp*) was weak, this synapse did not drive the spiking of the model (Figure 4C, up left). However, when *wSyn_supp* was strong, this synapse drove the spiking of the model (Figure 4C, down left). We made 100 trials for each *wSyn_supp* and at each trial the onset time and amplitude of the others synapses were chosen randomly to introduce spiking variability. We down-sampled our signal to 3 kHz and binarized it to 0 and 1, depending on the presence of APs or not similarly to (London et al., 2002). After calculating the information rate transfer using the quadratic extrapolations method, we obtained a sigmoid curve similar to previously published results (London et al., 2002) (Figure 4C, middle). To calculate the PNG Rate on binarized signals, we first converted our voltage signals to a 1-bit PNG image (2 levels of grey). As our signals are 100 trials of 5s at 3 kHz sampling, this yielded an image of 15 000 * 100 (= 1 500 000) pixels of 2 possible values. Similar to what has been described in paragraph 3.3, we obtained the PNG Rate of the signal measuring the size of the PNG file, dividing it by the number of pixels and multiplying it by the sampling to obtain a value in Bytes/s. To obtain the PNG entropy rate of the noise, we rotated the image 90 degrees, measured the size of this new file, divided it by the number of pixels and multiplied it by the sampling to obtain a value in Bytes/s. As expected, the difference between the PNG Rate of the signal and the PNG Rate of the noise followed a sigmoid curve similar to the one calculated by the quadratic extrapolations method (Figure 4C, right).

From this, we concluded that by saving multiple trials of the same experiment as a single PNG file, we can estimate the PNG Rate of the signal. And by simply rotating this same file 90 degrees and saving it again, we can estimate the PNG Rate of the noise. The difference between those two values follows the same behavior as measuring the information transfer rate between the stimulation protocol and the multiple recorded responses. Therefore, PNG Rate method is suitable to compare the Synaptic Information Efficacy of synapses displaying various strengths.

### Application to Place Fields

Some hippocampal cells have their firing rate modulated by the animal position, discharging specifically at a spatial region known as the place field of the cell (O’Keefe and Dostrovsky, 1971). Properly identifying these cells requires estimates of the information contained in spikes about navigational features (i.e., position, speed, head angle). The main metrics used to estimate this type of information were proposed by (Skaggs et al., 1993) and are derivations from Shannon’s mutual information. By definition, the experiment conditions are less controlled than in our previous simulations, as the experimenter cannot be certain that the animal will explore its full environment.

We generated a set of data mimicking an animal running randomly in an enclosed environment (Figure 5A). This closed environment was divided in 20*20 bins of space; the mock animal was put at a random point in the environment and was let to stay in for different durations. Animal speed and occupancy were considered constant over time and of 1 second per bin. We defined a simple “random walk” algorithm where at each time, the next position of the animal was chosen randomly between the 8 adjacent space bins and its own bin. This gives a random pathway exploring only some part of the environment, passing through the same space bin multiple times, etc… At each step, white noise of 1 spike was emitted with a chance of 0.1. In addition we set a mock place field, where until 20 spikes could be emitted following a Gaussian-like centered in the middle of the environment (width of ∼5 bins). This place field had an increasing chance to fire (from 0 to 1), in order to mimic place fields of increasing strengths. This gave a crude but versatile place field simulation, where we could not control the path of the mock animal (Figure 5A).

To estimate the spatial information contained in the firing rate of each cell, we computed the *I*_*sec*_ metric as described in (Skaggs et al., 1993; Souza et al., 2018), as it is equivalent to the Mutual Information from the average firing rate (over trials) in the N space bins using the following definition:

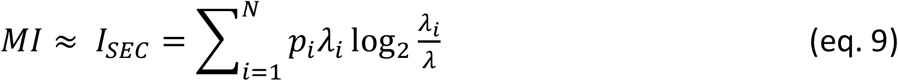

Where ⍰_*i*_ is the mean firing rate in the i-th space bin and p_i_ the occupancy ratio of the bin, while the overall mean firing rate of the cell. *I*_*sec*_ measures information rate in bits per second (Skaggs et al., 1993). This way of computing MI looks extremely simple, but by using it the user makes four essential assumptions: **i**) The information is purely encoded by the spike frequency; **ii**) The position of the animal is the only parameter which could influence the spike frequency; **iii**) The binning in time and space is ideal and **iv**) The spiking frequency of the noise and the place field will be similar between animals (But see (Souza et al., 2018) for an attempt to correct for this bias). By doing so, we can simplify our entropy and MI calculations to a single probability space and obtain equation 9. The direct methods (such as the quadratic extrapolations method) and the source compression methods (such as the PNG Rate) are generic ways of calculating entropy, without making any assumptions. However, in these simulated data we know that the position will be the only parameter affecting the spiking frequency, and thus the PNG Rate method should give equivalent results to equation 9.

We ran 10 simulations of 10 000 steps for each value of place field strength and calculated the MI according to (eq. 9). We obtained an increase of the MI with the strength of the place field (Figure 5C, left), as expected. To use our PNG algorithm, we simply traversed all the space bins with a pairing function, accumulating the spike values if there was one (Figure 5B). Briefly, we constructed a table with in x-axis the position bins of the image and in y-axis the number of times the animal passed into a given bin (Figure 5B, right). The value inside the case (i,j) of the table is the number of spikes emitted by the cell at the j^th^ time the animal passed into the i^th^ bin position (color from blue to red in Figure 5B, right). This gave us a 2D image with a 20*20 = 440 maximum width and a height depending on the number of times the mock animal went on the same position (Figure 5B, right). There were “holes” as it never went through every position the same amount of time, but we considered them as empty with 0 spike (grey case in Figure 5B, middle). We saved those images in both orientations, calculated the PNG rates and subtracted one to the other. The obtained curve has similar characteristics to the previously calculated MI (Figure 5C, right).

From this third test, we conclude that we can apply the same algorithm to more complex data like Place Fields recordings. Even if the data are not homogenous, a simple pairing transform can map all 2D coordinates to unique 1D index, progressively building a 2D image (Figure 5B). This image can then be saved as PNG to estimate its Entropy Rate and Mutual Information.

### Application to histology

Another way to understand entropy is that it is a representation of complexity of a signal (Cover and Thomas, 2006). Shannon entropy is linear by design as the original work was about coding messages through a communication line. However, multiple attempts have been made to generalize it to 2D signals (Azami et al., 2019; Larkin, 2016; Sparavigna, 2019) and for example, (Gavrilov et al., 2018) used shearlet transformation (Brazhe, 2018) to characterize entropy and complexity in two-dimensional pictures of astrocytic processes. The PNG format has been used with the same idea (Wagstaff and Corsetti, 2010) to evaluate the complexity of biogenic and abiogenic stromatolites.

In the same spirit, we used the ddAC Neuron example from the FIJI distribution of the ImageJ software (Schindelin et al., 2012). This reconstructed drosophila neuron (Figure 6Ba) is a classic example used for Sholl analysis (Ferreira et al., 2014; Sholl, 1953), (see https://imagej.net/Sholl_Analysis). This analysis estimates the complexity of an arborization by drawing concentric circles centered on the soma of the neuron and counting the number of intersections between those circles and the dendrites. The more intersections, the more complex is the dendritic tree. We realized a cylindrical anamorphosis centered on the soma of the ddAC neuron (Figure 6Bb) and saved each column of this new rectangular image as PNG files. As a result, the size of those files grew with the distance from soma, reaching the same peak as a Sholl analysis made with default settings in Fiji (Figure 6Bc). Of course, it is also possible to simply tile the original image in smaller PNG files and save them independently. The size of these files will give an idea of the complexity of the area covered by the tile (Figure 6Bd).

**Figure 6:**
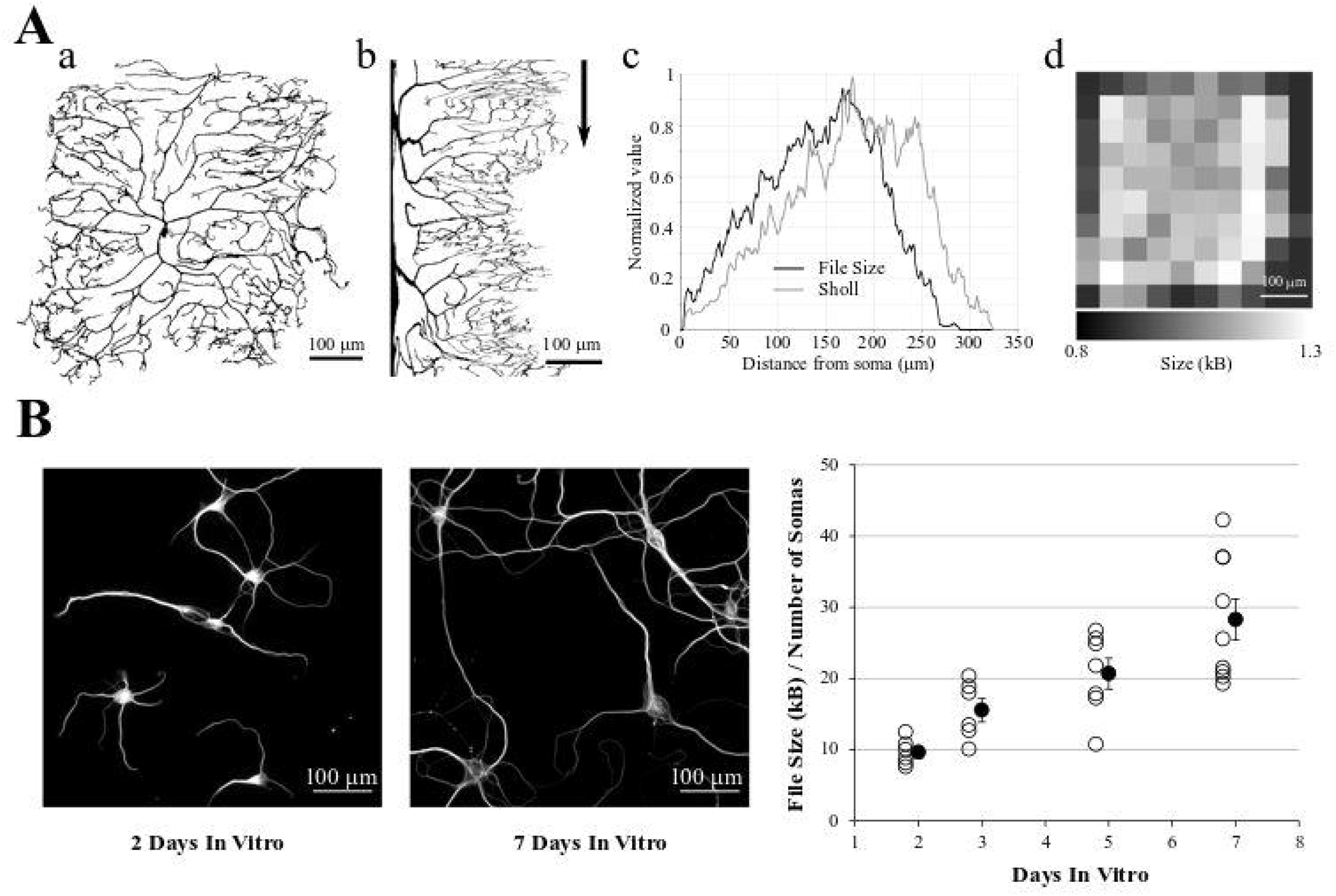
Application to 2D data. **A)** a: ddAC Neuron from Fiji examples. b: The same neuron after a circular anamorphosis centered on the soma. Note how the complexity of the dendritic arbor changes versus distance from soma. Arrow: each column of pixel was saved as a single PNG file. c: file size of these columns as PNG, showing the growth in complexity of the dendritic arborization (black line). As a comparison, we performed a Sholl analysis of the same image with default FIJI parameters (grey line). d: the original image was tiled in 10 squares, and each square saved as a PNG file. The sizes of these files reveal the heterogeneity of the dendritic arborization. **B)** Left and Middle: examples of MAP2 stainings of Brandner and Withers neuronal cultures at 2 and 7 Days In Vitro. Note the growth in dendritic arborization through time. Right: Each image was saved into a PNG file, and the file size divided by the number of visible somas. This gives us a file size normalized by the density of the culture. This value increases with the number of Days In-Vitro, revealing the dendrite growth.

In a final example, we used a group of images made by Dieter Brandner and Ginger Withers available in the Cell Image Library (http://cellimagelibrary.org/groups/3006). These images are under Creative Common Attribution Licence and show the growth of neuronal cultures from 2 to 7 days In-Vitro. They show two stainings, for tubulin and MAP2. They are suitable to our needs as all the images have the same dimensions and resolution. We kept only the MAP2 channel as it reveals the dendrite morphology, converted the images to 8-bits grey scale (256 grey levels) and thresholded them to remove the background (Figure 6C, left and middle). We then saved all the images to PNG, measured the size of the files on the hard drive and divided this number by the number of visible somas, in order to make a quick normalization by the culture density. As expected, this ratio File Size / Number of Cells increases with the number of days in culture, revealing the dendrite growth (Figure 6C, right).

From this fourth test, we showed that we can use the PNG format to estimate the entropy of 2D images as well, and this can be used to estimate dendrite growth or local entropy of an image.

## Discussion

Entropy measurement can be a tool of choice in neuroscience, as it applies to many different types of data; it can capture nonlinear interactions, and is model independent. However, an accurate measure can be difficult as it is prone to a sampling bias depending of the size of the recorded signal, its quantization levels and its sampling. There are multiple ways to compensate for it, but none of them trivial. In this paper, we showed that it is possible to estimate the entropy rate of neuroscience data simply by compressing them in PNG format, measuring the size of the file on the hard drive and dividing by the number of pixels. We called this measure the PNG Rate. The principle relies on the Source Coding Theorem specifying that one cannot compress a signal more than its entropy rate multiplied by its size. We showed first that the PNG Rate correlates linearly with the calculated Entropy Rate of white-noise-like signals (Figure 1). Then, we showed that the PNG Rate needs a minimal amount of data points to converge to a stable value. This allows comparing different files of different sizes, as long as we reached this stable rate (Figure 2). Moreover, we showed that this PNG Rate method is suitable to replace methods used in previously published articles, such as the role of AP frequency in Entropy Rate ((de Polavieja et al., 2005), Figures 3 and 4) or the impact of synaptic strength on the postsynaptic firing ((London et al., 2002), Figure 4). Finally, we showed that this method is applicable to the detection of place cells or the estimation of the complexity of neuronal arborization (Figures 5 and 6).

### Drawbacks of the PNG method

The main drawback of this method is that the PNG Rate is not the absolute value of the entropy rate of the signal. Even if entropy bits and computer bytes do share similar names, in no cases should we exchange one for the other. The PNG Rate is a way to estimate the evolution of entropy, considering all other parameters unchanged. As so, PNG files must be of the same dynamic range and saved with the same software. A PNG file is composed of a header, critical chunks and non-essential ancillary chunks (See Methods). Different software will save different data in the ancillary chunks, may filter the signal before compressing it and thus will change the size of the file, independently of the compressed signal.

Other works described how to normalize the compressed chain in order to infer the original entropy value (Avinery et al., 2019). But the experimenter has to keep in mind that compression algorithms perform better with long data chains and convergence to entropy rate is slow (see (Baronchelli et al., 2005; Benedetto et al., 2002; Goodman, 2002; Khmelev and Teahan, 2003) for an extensive discussion about the benefits and drawbacks of compression algorithms). We show here that as long as we have enough pixels in the generated images, the PNG Rate will converge toward a stable value (Figures 2 and 3), which allows comparing recordings of different lengths. This PNG Rate value is not the Entropy Rate itself, but methods able to make comparisons between conditions are often required and this method fits in this situation.

### Advantages of the PNG method

The main advantage of this method is that it relies on previously developed compression algorithms that were already shown as optimal (Huffman, 1952; Ziv and Lempel, 1977). Moreover, it does not need any specialized software or any knowledge in programming language, as the PNG format is ubiquitous in informatics. For example, the ImageJ software is widely used in neuroscience and can export data as PNG.

A second advantage is the speed of execution. As an example, the information transfer rates calculation (Figure 4) took a bit more than 2 hours for the quadratic extrapolations method. Saving the same signals in PNG to calculate the PNG Rates took less than 30 seconds on the same laptop computer.

As so, this method is extremely easy, quick, and does not need any knowledge in mathematics for correcting the sampling bias. It is interesting to note that an experimenter will often acquire multiple recordings of the same protocol in order to infer proper statistics. This means that most of the times no supplementary experiments are needed to calculate the entropy rate of a signal, or the information transfer rate between a stimulation protocol and its recorded result.

In conclusion, we propose the PNG method as a quick-and-easy way to estimate the entropy rate of a signal or the information transfer rate between stimulation and recorded signals. It does not give the exact value of entropy rate or information transfer rate, but it is related to these values in a linear way which allows the evaluation of their evolution in different experimental conditions.

### When to use it?

As shown in this study, the PNG Rate method is simple and can be used as a back-to-the-envelope way to measure any change in entropy rate or information transfer rate through different experimental conditions. It does not give the absolute value of entropy, but often the experimenter wants a simple comparison to a control situation. This method can be virtually used on every kind of neuroscientific data. Here, we showed examples of applications on patch-clamp data, detection of place cells by extracellular spiking recordings or histological data. Moreover, if the experimenter needs to compare multiple files of different lengths, it is possible to calculate the PNG Rate in portions of the signal with different sizes and find the minimal number of points needed for the PNG Rate to stabilize. If the number of points in all experimental conditions is above this minimal number, their PNG Rates can be compared to estimate the variation of entropy rate in the different conditions. Several works used compression algorithms to estimate absolute value of entropy. However, we do not think they apply to our simple method. The first method needs to have access to the dictionary created by the compression algorithm (Baronchelli et al., 2005). However, with the PNG algorithm we do not have access to the dictionary itself. The second method needs to first estimate the maximum entropy possible by the model (Avinery et al., 2019), which is possible as well but will need more conventional algorithm to be determined.

### Developments

We see multiple ways to improve this method. First, we saved our data as 8-bits PNG files, which limits the dynamic range of the file to 256 values. However, it is possible to save PNG natively as 1, 4, 8, 16 and 24 bits range, thus greatly increasing the dynamic range of the saved signal. Second, with some programming skills it is possible to remove the header and ancillary chunks of the PNG format, thus removing the size overhead (but the file will be unreadable by standard softwares). Finally, one possible way of improving the estimation of entropy rate would be to choose a better compression algorithm. We choose the PNG format as it is widely used by common softwares and it is based on LZSS and Huffman algorithms, which have been proven optimal. However, some algorithms may give a better compression rate depending on the quality of the data. As an example, the Rice compression algorithm was originally developed for the NASA Voyager missions (Rice and Plaunt, 1971). It is suboptimal but is better suited for noisy signals of low values.

In a more general direction, it is important to note that this method works with any entropy-coding compression algorithm, as long as they are loss-less. This is the case of GZip algorithms for example, used in many compression softwares such as WinRAR, PKZIP, ARJ, etc… It is thus not limited to pictures in PNG, although this format is useful for rotating the file and estimating the mutual information easily. Moreover, we apply these algorithms to 2D images, when actually the algorithm linearizes the data and works only in linear way on one dimension. There are some attempts to generalize Shannon entropy to 2D space (Azami et al., 2019; Brazhe, 2018; Larkin, 2016; Sparavigna, 2019), but they are out of the scope of this paper.

## Data accessibility

Script and codes are available online: https://github.com/Sylvain-Deposit/PNG-Entropy

## Supplementary material

Script and codes are available online: https://github.com/Sylvain-Deposit/PNG-Entropy

## Acknowledgements

We thank Q. Bernard, C. Leterrier, V. Marra, T. Jensen, K. Zheng, S. Martiniani, R. Beck for helpful discussions. Two first versions of this manuscript have been released on bioRxiv (Zbili and Rama, 2020), https://www.biorxiv.org/content/10.1101/2020.08.04.236174v2

Version 3 of this preprint has been peer-reviewed and recommended by Peer Community In Circuit Neuroscience (https://doi.org/10.24072/pci.cneuro.100001).

## Conflict of interest disclosure

The authors declare that the research was conducted in the absence of any commercial or financial relationships that could be construed as a potential conflict of interest.

Sylvain Rama is a recommender for PCI Circuit Neuroscience.

